# CADD from *Chlamydia trachomatis* is a manganese-dependent oxygenase that employs a self-sacrificing reaction for the synthesis of *p*-aminobenzoate

**DOI:** 10.1101/2022.08.01.502304

**Authors:** Rowan Wooldridge, Spenser Stone, Andrew Pedraza, W. Keith Ray, Richard F. Helm, Kylie D. Allen

## Abstract

CADD (<u>c</u>hlamydia protein <u>a</u>ssociating with <u>d</u>eath <u>d</u>omains) is a *p*-aminobenzoate synthase involved in a non-canonical route for tetrahydrofolate biosynthesis in the intracellular bacterial pathogen, *Chlamydia trachomatis*. The previously solved crystal structure revealed a seven-helix bundle architecture similar to heme oxygenase with a diiron active site, making CADD a founding member of the emerging HDO (<u>h</u>eme-oxygenase-like <u>d</u>iiron <u>o</u>xidase) superfamily. The CADD-dependent route for pAB biosynthesis was shown to use L-tyrosine as a precursor, however, *in vitro* reactions were not stimulated by the addition of free L-tyrosine or other tyrosine-derived metabolites, leading to the proposal that the enzyme uses an internal active site tyrosine residue as a precursor to pAB. Here, we perform further biochemical characterization of CADD to clarify the details of the unique self-sacrificing reaction. Surprisingly, the pAB synthase reaction was shown to be dependent on manganese as opposed to iron and the data are most consistent with an active dimanganese cofactor analogous to class Ib and class Id ribonucleotide reductases. Experiments with ^18^O_2_ demonstrated the incorporation of two oxygen atoms from molecular oxygen into the pAB product, supporting the proposed mechanism requiring two monooxygenase reactions. Mass spectrometry-based proteomic analyses of CADD-derived peptides demonstrated a glycine substitution at Tyr27, a modification that was increased in reactions that produced pAB *in vitro*. Additionally, Lys152 was found to be deaminated and oxidized to aminoadipic acid. Taken together, our results support the conclusion that CADD is a manganese-dependent oxygenase that uses Tyr27 and possibly Lys152 as sacrificial substrates for pAB biosynthesis.

## Introduction

Tetrahydrofolate (THF, Figure 1A) and its derivatives – collectively referred to as folates – are essential cofactors for one-carbon transfer reactions in all domains of life. Animals are dependent on their diet for folate acquisition; however, most bacteria, plants, and fungi synthesize folates *de novo. Chlamydia trachomatis* is an intracellular bacterial pathogen that utilizes several non-canonical enzymes for THF biosynthesis (1,2). The *p*-aminobenzoate (pAB) portion of THF is normally synthesized via chorismate by two enzymes – the bifunctional aminodeoxychorismate synthase (encoded by *pabA* and *pabB*) and aminodeoxychorismate lyase (encoded by *pabC*). However, *C. trachomatis* is missing these three genes and instead uses the product of a single gene, *ct610*, in a route that does not directly employ chorismate (2). Before the discovery of its role in THF biosynthesis, the *ct610* gene product was named CADD (<u>c</u>hlamydia <u>p</u>rotein <u>a</u>ssociating with <u>d</u>eath <u>d</u>omains) due to its role in inducing host cell apoptosis via interacting with death domains of mammalian tumor necrosis factor family receptors (3). Thus, the *ct610* gene product appears to be a “moonlighting” protein with at least two distinct and seemingly unrelated functions. The crystal structure of CADD revealed a dimer of seven-helix bundles (4) (Figure S1A), an architecture first observed in heme oxygenase (5), with an active site containing a diiron cofactor (Figure 1B and Figure S1B) similar to the hydroxylase component of soluble methane monooxygenase (sMMOH) and the β subunit of class Ia ribonucleotide reductase (RNR) (4). Thus, CADD is a member of the emerging <u>h</u>eme-oxygenase-like <u>d</u>iiron <u>o</u>xidase (HDO) superfamily, of which ~10,000 members are bioinformatically proposed, but only a handful have defined enzymatic activities (6). Characterized HDOs catalyze diverse reactions including *N*-oxygenation (SznF (6–9), RohS (10), and FlcE (11)), oxidative C-C bond cleavage (UndA (12–14), BesC (15–17), and FlcE (11)), and methylene excision (FlcD) (11).

**Figure 1.**
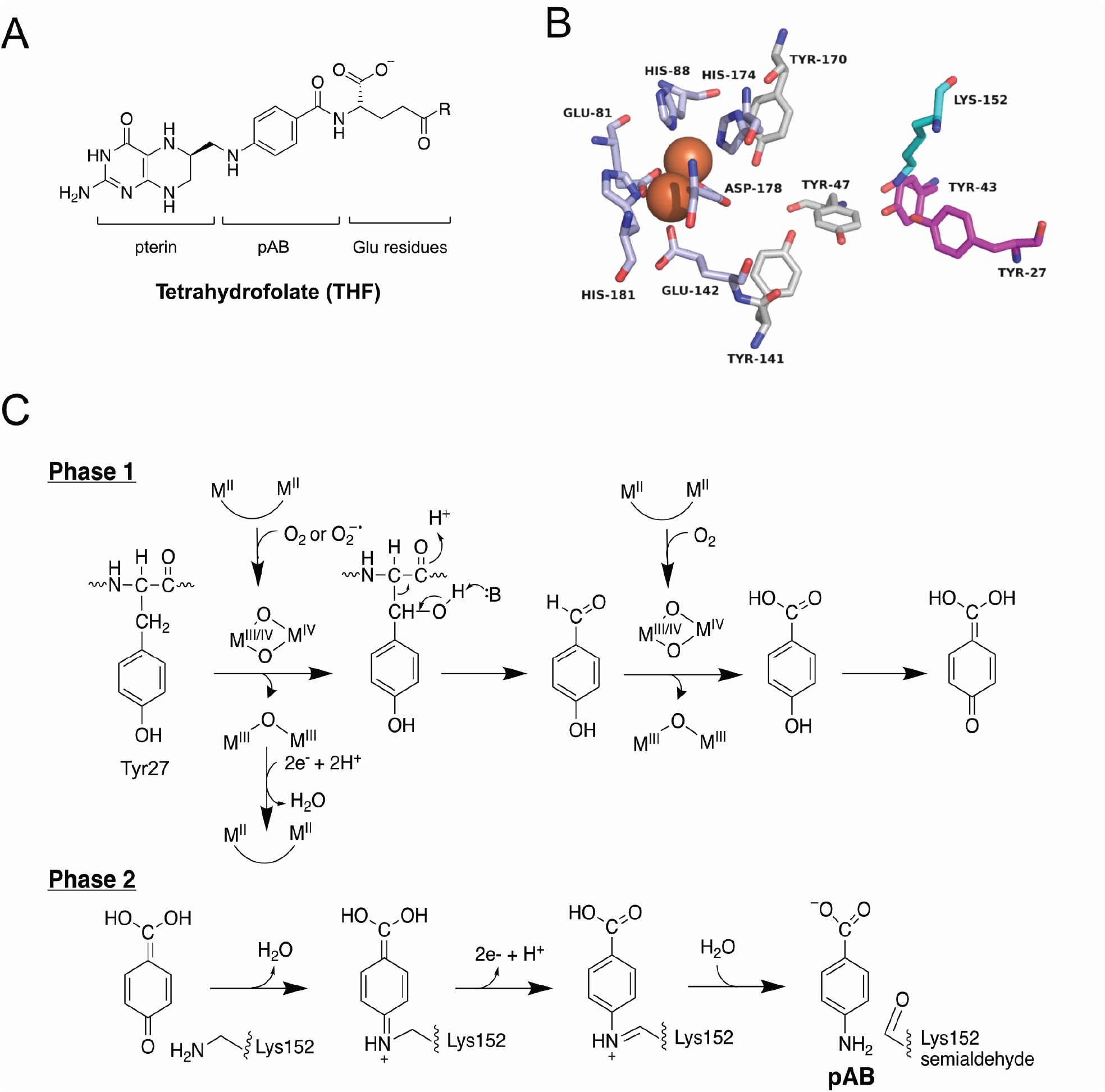
CADD’s proposed involvement in pAB biosynthesis. (A) structure of THF; (B) active site of CADD (pdb ID: 1RCW) with residues of interest highlighted - diiron-ligating residues are shown in light blue, Tyr residues previously shown to be essential for pAB synthase activity are shown in magenta while Tyr residues shown to be non-essential for pAB synthase activity are in depicted grey; Lys152 is shown in cyan. (C) proposed self-sacrificing mechanism for pAB synthesis. “M” represents a metal ion.

Although CADD was the first HDO superfamily member with a reported crystal structure (4), its enzymatic activity(ies) and physiological function(s) remain unclear. We previously utilized isotope feeding studies in an *E. coli* ΔpabA strain expressing CADD to demonstrate that L-tyrosine is the likely substrate for CADD-dependent pAB biosynthesis (18). Further, *in vitro* studies demonstrated that the purified enzyme produces pAB in a reaction that requires a reducing agent and molecular oxygen, but does not appear to utilize free L-tyrosine as a substrate. Site-directed mutagenesis identified two active site tyrosine residues (Tyr27 and Tyr43, Figure 1B) as being essential for pAB synthesis and, thus, we suggested that CADD is a self-sacrificing enzyme that cleaves an active site tyrosine residue from the protein backbone to serve as a substrate for pAB biosynthesis (18). Additionally, mutating a conserved lysine residue (Lys152) to arginine resulted in a dramatic decrease in pAB production, suggesting it could be the missing amino group donor in the reaction. This sacrificial use of a lysine residue as an amino group donor has been reported for BioU, an enzyme involved in an unusual route for biotin biosynthesis in cyanobacteria (19).

In our proposed working mechanism, radical-mediated hydroxylation employing an oxygenated-dimetal cofactor produces a β-hydroxylated tyrosine residue. This activates the residue for acid-base catalyzed cleavage from the protein backbone to generate *p*-hydroxybenzaldehyde, followed by another monooxygenase reaction to produce *p*-hydroxybenzoate (pHB) (Figure 1C, phase 1). The amination occurring in phase 2 of the reaction is inspired by the BioU reaction (19) as well as copper-dependent amine oxidases that use tyrosine-derived quinone cofactors (20). Thus, the pHB intermediate could tautomerize to form a quinone methide species that reacts with the ε-amino group of Lys152 to produce a Schiff base intermediate. Oxidation to the product-like Schiff base followed by hydrolysis completes the reaction to yield Lys152-semialdehyde and the final pAB product (Figure 1C, phase 2). Here, we further characterize this enigmatic pAB synthase to clarify its metal-dependence and elucidate key details of the proposed reaction.

## Results

### Metal-dependency of pAB synthase activity of CADD

CADD containing a 10X-His-tag was heterologously expressed and purified from *E. coli* (Figure S2) (18). As-purified, CADD contained ~0.05-0.1 Fe ions per monomer, indicating that the majority of the enzyme population lacks the expected diiron cofactor. This result is consistent with other HDO family members (6,7,9,12,13,17), where the metallocofactor is apparently highly unstable. LC-MS analysis of reactions containing as-purified CADD (154 µM) under aerobic conditions showed the production of ~3 µM pAB when the enzyme was incubated with dithiothreitol (DTT) versus ~1 µM pAB detected when incubated without a reducing agent (Figure 2A). As previously reported (18), the addition of possible small-molecule substrates (L-tyrosine, *p*-hydroxyphenylpyruvate, pHB, or chorismate) does not increase pAB production, suggesting that the enzyme uses a protein-derived substrate or is purified with an unknown substrate bound. The DTT is expected to be responsible for generating the reduced metal center that can react with molecular oxygen (or a reactive oxygen species) to produce a peroxo species involved in the two proposed monooxygenase reactions (Figure 1C). The pAB present in the reaction without DTT is likely due to minor amounts of pAB from the *E. coli* expression host remaining bound to the purified enzyme. Several other reducing agents were tested, but DTT resulted in substantially more pAB production compared to other chemical reductants (Figure S3). Oxygraph analysis showed that as-purified CADD consumed oxygen at a rate of ~0.12 min^-1^ when the enzyme was reduced with DTT and virtually no oxygen consumption was detected when a reducing agent was not included in the reaction (Figure 2B and 2D).

**Figure 2.**
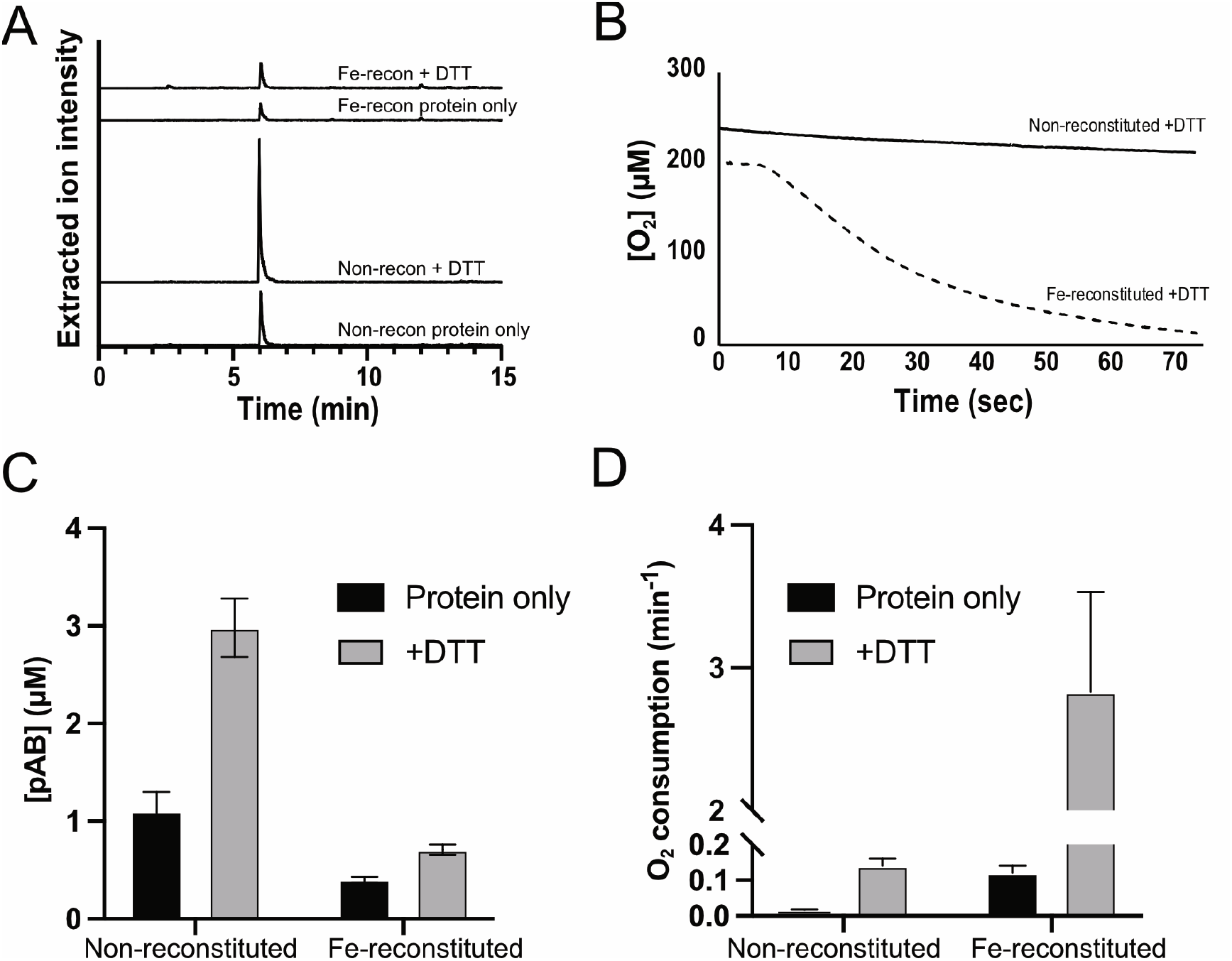
pAB synthase and oxygen consumption activities of non-reconstituted compared to Fe-reconstituted CADD. (A) LC-MS analysis of pAB produced in different CADD reactions, chromatograms represent an extracted ion current corresponding to two pairs from a multiple reaction monitoring method specific for pAB. (B) Oxygraph analysis of oxygen consumption by CADD. (C) Quantitation of pAB produced by non-reconstituted vs. Fe-reconstituted CADD. (D) Quantitation of O_2_ consumption by non-reconstituted vs. Fe-reconstituted CADD. Error bars represent the standard deviation of at least three replicates.

The exceedingly low Fe content in the as-purified enzyme was expected to result in the very low *in vitro* pAB synthase activity. Thus, we reconstituted the enzyme in the presence of 2x molar excess Fe(II) under anaerobic conditions. This resulted in nearly complete incorporation of a diiron cofactor (1.75 ± 0.3mol Fe per mol monomer) as measured by a ferrozine iron determination assay. Surprisingly, however, Fe-reconstituted CADD exhibited significantly decreased pAB synthase activity compared to the non-reconstituted enzyme (Figure 2A and 2C). In contrast, the rate of oxygen consumption of the Fe-reconstituted protein increased from ~0.12 min^-1^ to ~3.0 min^-1^ (Figure 2B and 2D). These results suggest that the production of an oxygenated diiron cofactor is not coupled to the pAB synthase activity, thus leading us to hypothesize that an alternative metal cofactor was required.

To test this, CADD was reconstituted with Zn(II), Cu(II) and Mn(II) as well as Fe(II) paired with each of these three metals. Our initial experiments using equimolar Fe with other metals resulted in very low pAB synthase activity, likely due to the formation of Fe/Fe cofactors. Thus, in subsequent di-metal reconstitutions, a sub-stochiometric amount of Fe was added (0.5x) to mitigate the formation of Fe/Fe cofactors as opposed to the desired heterodimetallic species. The presence of all three alternative metals produced an increase in pAB synthase activity compared to the non-reconstituted and Fe-reconstituted enzyme, with Mn showing the largest increase in pAB production (Figure 3A). Interestingly, the Mn-only and Mn/Fe reconstitutions resulted in similar pAB synthase activity in this experiment.

**Figure 3.**
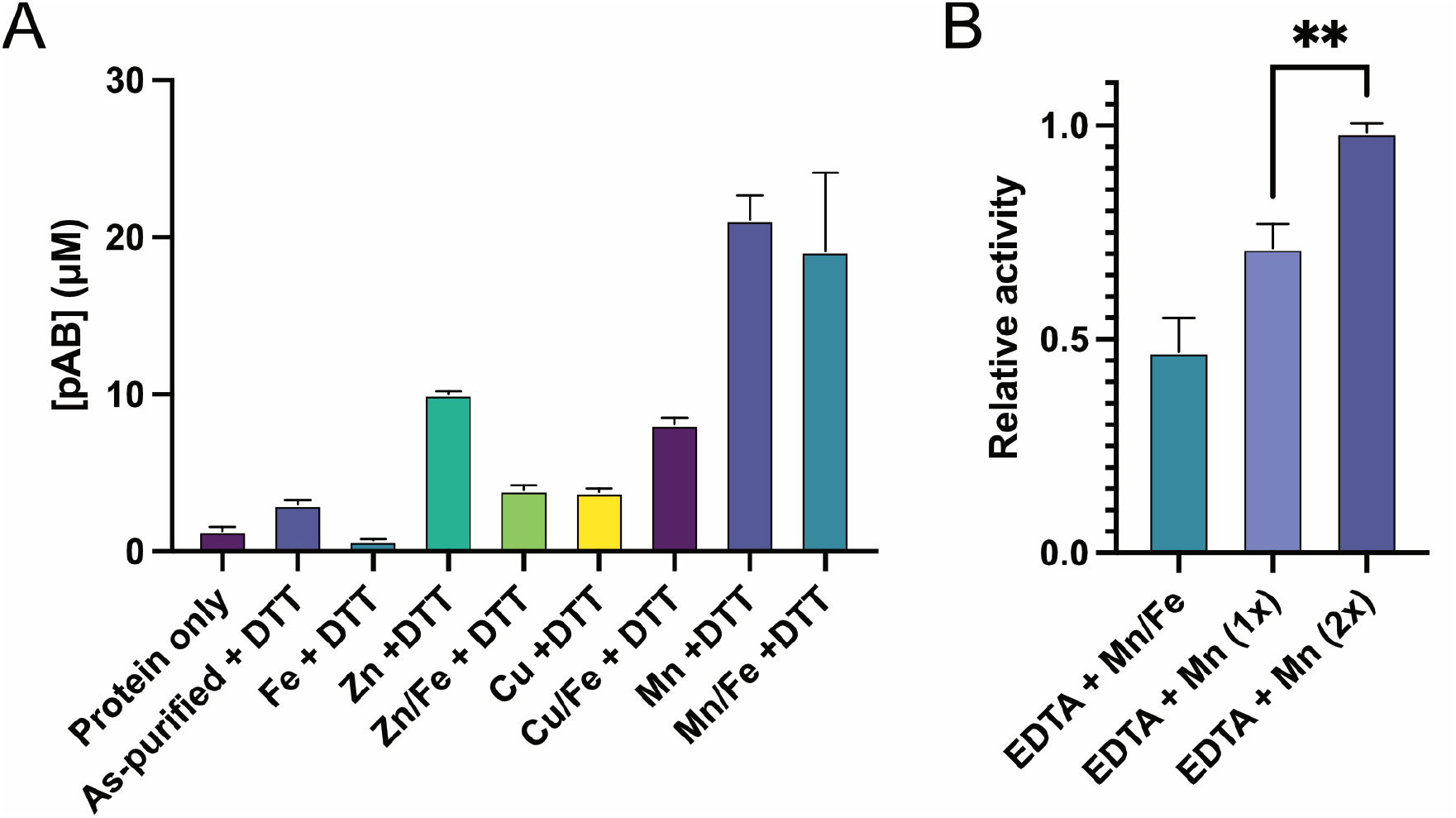
Metal-dependence of CADD pAB synthase activity. (A) pAB produced in assays with CADD reconstituted with various metals, then incubated with DTT under aerobic conditions. (B) pAB synthase activity of EDTA-treated CADD reconstituted with indicated metals and incubated with DTT under aerobic conditions. Error bars represent the standard deviation of at least three different replicates. Statistical analysis of the 1x vs. 2x Mn was performed using a Welch’s t-test in Prism 9. **p-value = 0.0075.

To confirm the identity of the metal cofactor required for pAB synthesis, as-purified CADD was treated with EDTA to remove any existing metals bound and then the apo-protein was reconstituted with Fe or Mn, followed by assaying for pAB synthase activity. The Mn-reconstituted protein showed ~10-fold higher pAB synthase activity compared to the Fe-reconstituted protein (Figure S4), thus confirming that the pAB synthase activity of CADD is dependent on Mn and not Fe. Since the initial Mn/Fe and Mn-only experiments revealed similar activities (Figure 3A), we performed additional EDTA experiments to clarify the metal-dependence of pAB synthesis by CADD. Reactions with 2x-Mn added showed the highest activity, while both 1x Mn and 1xMn/0.5x Fe produced significantly less pAB (Figure 3B).

Taken together, our current data are most consistent with the pAB synthase activity of CADD being solely Mn-dependent. Given the increase in activity when supplemented with 2x Mn vs. 1x Mn (Figure 3B) and due to the established identity of dimanganese active sites in related class 1b and class 1d RNRs, the active cofactor for the pAB synthase activity of CADD is most likely a dimanganese species as opposed to a mononuclear manganese site.

In addition to monitoring pAB synthase activity, we tested the oxygen reactivity of various forms of metal-reconstituted CADD. None of the reconstitutions showed a significant increase in the rate of oxygen consumption like observed in the Fe-reconstitution (Figure S5). An increase in oxygen consumption was seen primarily in the mixed reconstitutions containing Fe, likely due to an increase in the amount of CADD containing a Fe/Fe cofactor, which is most reactive towards O_2_. Since, the oxygenated diiron cofactor did not seem to be used for pAB production, we assayed for the production of H_2_O_2_ as a potential unproductive product of the Fe-reconstituted CADD. We found that Fe-reconstituted CADD produced high levels of H_2_O_2_ in the presence of DTT and O_2_ (~40 µM produced in 30 s) compared to Mn-reconstituted CADD that produced levels of H_2_O_2_ similar to controls lacking enzyme (Figure S6). Taken together, our data indicate that Fe-reconstituted CADD reacts with O_2_ to produce an oxygenated diiron cofactor, but this is not coupled to pAB synthesis and instead forms H_2_O_2_ as a side-product.

Given that the pAB synthase activity of CADD likely involves a dimanganese cofactor, we next considered how the active oxygenated cofactor would be generated. The dimanganese sites of previously characterized RNRs are unreactive towards O_2_ itself and require superoxide 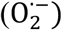 instead to generate the active cofactor. In the case of class Ib RNRs, a specific activase enzyme (NrdI) is used, while in class Id RNRs, no activase protein has yet been identified but the *in vitro* reaction is stimulated by the addition of superoxide-generating compounds such as hydroquinol (HQ) (21,22). Thus, we tested the activity of Mn-reconstituted CADD in the presence of HQ compared to a control sample from the same protein prep that lacked HQ. We observed a modest increase in pAB synthase activity in the presence of HQ at early timepoints in the reaction (0.73 µM/min vs. 0.52 µM/min without HQ). However, after 1 hr, the reaction without HQ produced more pAB compared to the reaction with HQ (Figure S7). Thus, the formation of the oxygenated-dimanganese cofactor is likely not the major rate-limiting step for pAB formation in our *in vitro* reactions and the amount of superoxide naturally present in the aqueous solution without HQ is enough to support activity.

### Incorporation of molecular oxygen into pAB produced by CADD

We previously showed that O_2_ was required for the pAB synthase activity of CADD (18). To further probe the role of oxygen in the reaction, we carried out assays in the presence of isotopically-labeled O_2_ with Mn-reconstituted CADD. The pAB produced by CADD exposed to atmospheric ^16^O_2_ displayed a [M + H]^+^ ion at 138 m/z (Figure 4A), while the pAB produced by CADD in a sealed vial with provided ^18^O_2_ displayed a predominant [M + H]^+^ ion at 142 m/z (Figure 4B). The peak at 140 *m/z* is also enriched in this sample due to a mixed species with one ^18^O incorporated and one ^16^O incorporated as a result of minor contamination from atmospheric oxygen. This demonstrates that two oxygen atoms from O_2_ are incorporated into the pAB product, supporting our proposed mechanism where CADD utilizes two monooxygenase reactions to produce the carboxylic acid portion of pAB (Figure 1C, part 1).

**Figure 4.**
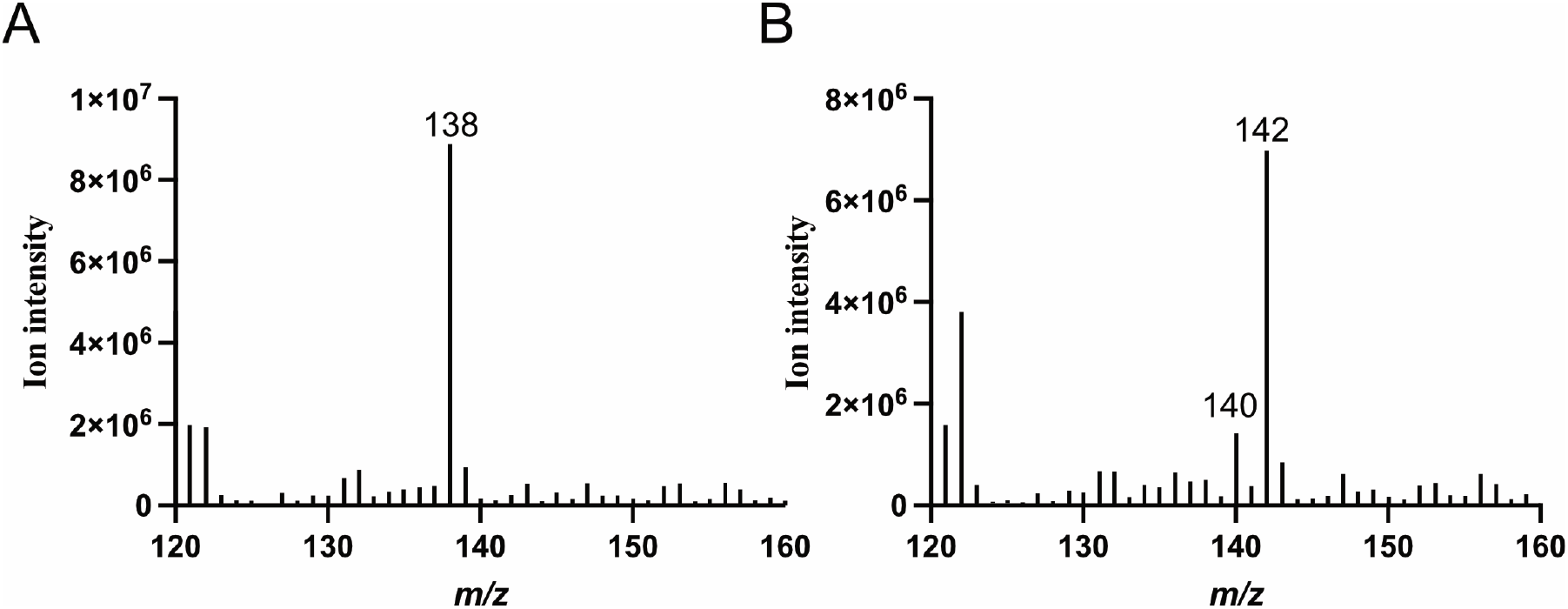
Incorporation of molecular oxygen into pAB by CADD. (A) pAB peak from Mn-reconstituted CADD reaction in the presence of atmospheric ^16^O_2_. (B) pAB peak from Mn-reconstituted CADD reaction in the presence of ^18^O_2_.

### Identification of sacrificial residues for pAB synthesis by CADD

One of the major questions regarding the pAB synthase activity of CADD is the identity of the substrate(s) for the reaction. Based upon feeding studies and preliminary enzymatic experiments, we previously proposed a self-sacrificing reaction mechanism (18), where the aromatic portion of a tyrosine residue is utilized as the substrate for pAB formation (Figure 1C, part 1). To provide evidence for this hypothesis, peptides resulting from protease-digested CADD were subjected to LC-MS analysis. Intriguingly, Tyr-27 was found to be converted to glycine in some HTFY_27_VKWSKGE peptides (monoisotopic mass for unmodified peptide is 1380.672 compared to 1274.640 for Tyr to Gly modification). The intensity of the ion corresponding to the Tyr27 to Gly modification increased ~50-fold in samples from *in vitro* reactions with Mn-reconstituted CADD in the presence of DTT compared to the absence of DTT (Figure 5B), thus providing strong support that this modification is linked to pAB formation (Figure 1C, part 1). The MS/MS spectrum of the [M + 2H]^+2^ ion shows characteristic fragment ions consistent with the identity of the Tyr27 to Gly modification (Figure 5C). Further analysis of CADD-derived peptides revealed that the Lys152-containing peptide, K_152_IRGLTE, also contains a modification, where Lys152 is deaminated and oxidized to aminoadipic acid (Figure 5D and 5E). The ion corresponding to the deaminated lysine was found to be ~10X more intense in Mn-reconstituted reactions with DTT compared to without DTT (Figure 5D), thus correlating the pAB synthase activity with the lysine modification. We previously showed that this lysine residue played a critical role in the pAB synthase reaction via site-directed mutagenesis (18), so the finding of deaminated Lys152 suggests the potential use of this residue as an internal amino group donor (Figure 1C, part 2). Since Lys residues can undergo non-specific oxidative deamination, we checked for the presence of other modified lysines in CADD and did not see any evidence of modification, thus providing further support that Lys152 is used as an amino group donor.

**Figure 5.**
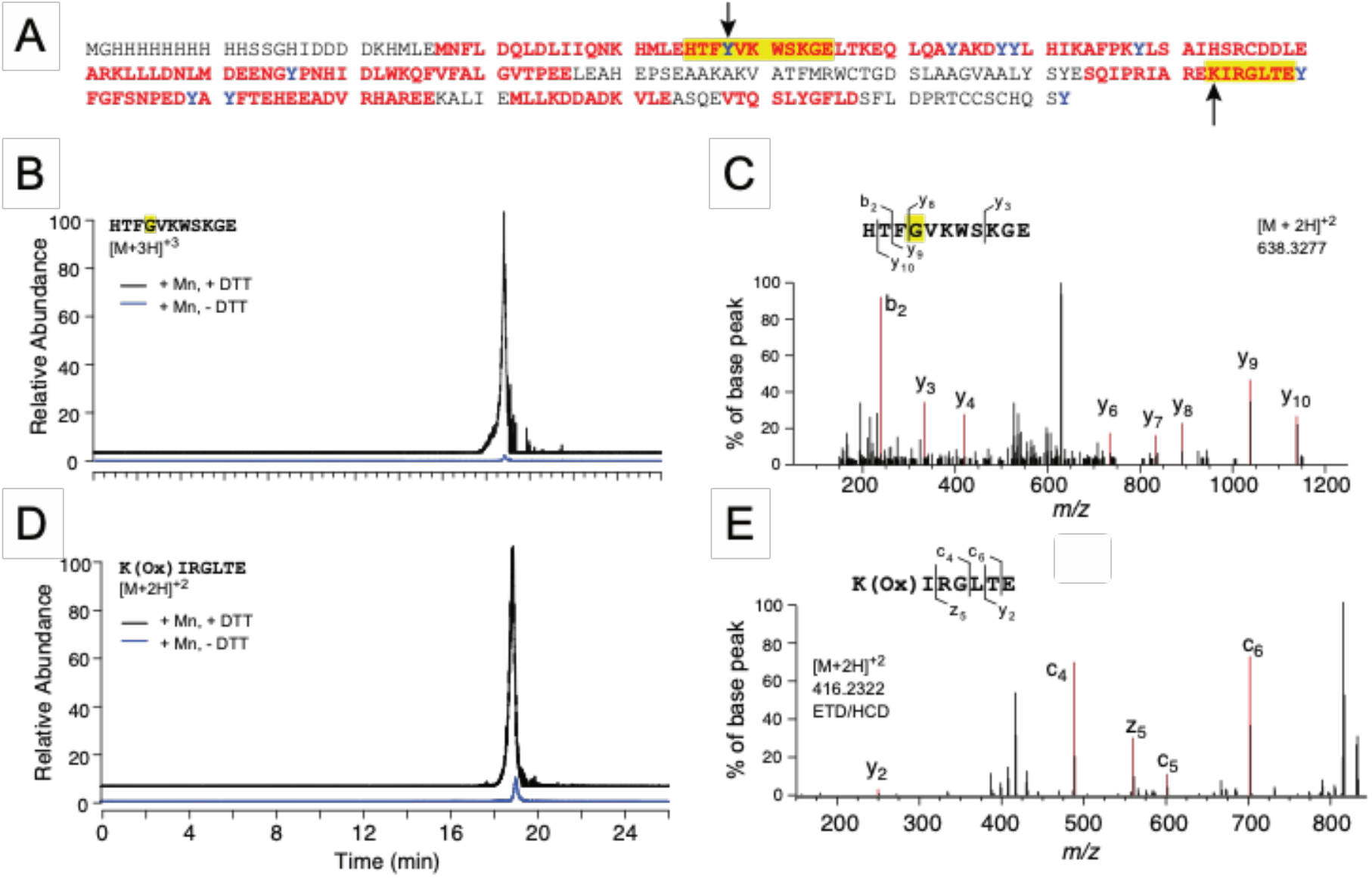
LC-MS analysis of modified amino acids in the CADD reaction. (A) Primary sequence of CADD expressed with a His-tag from pET19b. Peptides identified by LC-MS analysis of GluC-digested CADD are shown in red; peptides of interest containing Tyr27 and Lys152 are highlighted in yellow and black arrows indicate specific residues of interest. (B) Extracted ion chromatogram corresponding to the [M + 3H]^+3^ ion for the peptide containing Tyr27 to Gly modification in Mn-reconstituted CADD reactions with (black) and without (blue) DTT. (C) MS/MS spectrum of *m/z* = 638.3277, the [M + 2H]^+2^ ion, showing characteristic fragments for Tyr27 to Gly modification. (D) Extracted ion chromatogram corresponding to the [M + 2H]^+2^ ion for the peptide containing Lys152 to aminoadipic acid modification in Mn-reconstituted CADD reactions with (black) and without (blue) DTT. (E) ETD/HCD spectrum of *m/z* = 416.2322, showing fragments corresponding to the Lys152 to aminoadipic acid modification.

## Discussion

CADD is a founding HDO superfamily member involved in an unusual route for pAB biosynthesis in *C. trachomatis*. Based upon the crystal structure (4), which showed a diiron-containing active site (Figure 1B and Figure S1B), it was assumed that CADD would employ this diiron cofactor for pAB biosynthesis (18). However, here we showed that pAB formation by purified CADD is increased in the presence of manganese as opposed to iron. The use of Mn in place of Fe has established precedent in related and well-studied class I RNRs. A dimanganese active site is found in class Ib and Id RNRs, while class Ic RNR harbors a heterodinuclear Mn/Fe active site (22). In traditional class Ia RNRs, the reduced Fe^II^/Fe^II^ cofactor in the β subunit reacts with oxygen to generate a Fe^III/^Fe^IV^ species, which is used to generate a tyrosyl radical that undergoes long-range radical translocation to produce the catalytic cysteinyl radical in the α subunit (23). Interestingly, *C. trachomatis* utilizes a well-characterized class Ic RNR, where an oxygenated Mn^IV^Fe^III^ complex is the direct cysteinyl radical forming species (24) as opposed to the involvement of a tyrosyl radical.

Although RNR from *C. trachomatis* uses a Mn/Fe cofactor, our results indicate that the pAB synthase activity of CADD is solely Mn-dependent and currently disfavor the involvement of a Mn/Fe cofactor. This can be compared to class Ib and class Id RNRs, both of which employ an oxygenated Mn^III^/Mn^IV^ cofactor, where class Ib uses the cofactor to generate a tyrosyl radical and class Id utilizes the oxidized cofactor directly as the radical initiator (21,22,25). In both of these dimanganese enzymes, the reduced Mn^II^/Mn^II^ cofactor does not have sufficient reducing power to react with O_2_ directly (25,26), and instead requires the presence of superoxide for generation of the active oxidized cofactor (21,27). Superoxide occurs naturally in biological systems, but in the case of class 1b RNRs, this reactive species is specifically generated and delivered to the β subunit of class Ib RNR by a flavoprotein called NrdI (25,27,28). In the case of class Id enzymes, the *in vivo* source of superoxide is unclear, but superoxide-generating compounds such as HQ can generate the cofactor *in vitro* (21). Here, we found that HQ only slightly enhances the rate of pAB formation by CADD at early timepoints, thus indicating that the formation of the oxygenated-dimanganese cofactor may not be the primary rate-limiting step for pAB synthesis *in vitro* and the amount of superoxide naturally present in solution is enough to support activity. However, the true nature of the oxygen species that reacts with the dinuclear cofactor of CADD will require further detailed biochemical and spectroscopic studies to confirm.

The Mn-dependency of CADD as well as the use of Mn in the Mn/Fe cofactor of class Ic RNR in *C. trachomatis* as opposed to the canonical diiron cofactor of class Ia RNR may provide important insights into the unique metal biology of this intracellular pathogen. *C. trachomatis* is dependent on the availability of iron for growth, but iron homeostasis remains a poorly understood area of chlamydial biology (29). Importantly, successful pathogens must possess specialized mechanisms to overcome strict iron limitations imparted by the host (30). Thus, it is possible that *Chlamydiae* have adapted to use Mn in place of Fe in some cases. This is a strategy that has been observed in other pathogens such as the Lyme disease pathogen, *Borrelia burgdorferi*, which appears to not require iron at all and uses Mn instead of Fe for many traditionally Fe-dependent processes (31).

The canonical route for pAB biosynthesis uses chorismate as a substrate; however, genetic studies ruled out chorismate as a precursor for CADD-dependent pAB biosynthesis (2). Our previous feeding studies with CADD expressed in an *E. coli* ΔpabA strain showed that the aromatic portion of L-tyrosine was incorporated into pAB, but the addition of L-tyrosine to *in vitro* reactions did not result in enhanced pAB production (18). Additionally, site-directed mutagenesis revealed that two conserved tyrosine residues (Tyr27 and Tyr43) were essential for activity, leading us to hypothesize that CADD utilizes a self-sacrificing mechanism for pAB synthesis. Here, we showed that CADD contains a glycine substitution for Tyr27, a modification that was highly increased in reaction conditions associated with pAB production *in vitro*, thus strongly supporting the function of Tyr27 as a sacrificial substrate. Additionally, further analyses of CADD-derived peptides revealed the presence of 2-aminoadipic acid in place of Lys152. This result coupled with our previous data showing the critical role of this residue for pAB synthase activity (18) suggests that Lys152 may serve as a sacrificial amino group donor. However, these data must be interpreted carefully since lysine residues are known to be susceptible to oxidative damage via metal-catalyzed oxidation to generate 2-aminoadipic semialdehyde (lysine semialdehyde) (32), which is further oxidized to the more stable 2-aminoadipic acid (33). Thus, the source of the amino group donor for CADD-dependent pAB synthesis and the role of Lys152 will require further investigation.

Our current proposed working mechanism involves an initial monooxygenase reaction, where the *β* carbon of Tyr27 is hydroxylated to activate it for subsequent cleavage from the protein backbone, followed by a second monooxygenase reaction to generate the final carboxylic acid group (Figure 1C, part 1). In part 2 of the reaction, a hydroxyl group is converted to an amine facilitated by Schiff base chemistry with Lys152, where the amino group of this residue is eventually transferred to the substrate to generate the final pAB product. It is important to note that our current results do not provide any information about the order in which the tyrosine cleavage and amination reactions occur, so part 1 and part 2 in Figure 1C could be switched. A perplexing aspect of the proposed self-sacrificing reaction is that, based on the CADD crystal structure (4), the residues involved (Tyr27 and Lys152) are located ~13-14 Å away from the metallocofactor (Figure 1B). It is possible that a conformational change occurs upon reaction with oxygen that brings these residues closer to the oxygenated cofactor for the reaction to occur and/or that the active site of Mn-loaded vs. Fe-loaded CADD is reorganized, thus leading to pAB formation with Mn present, but not when Fe is present. Alternatively, the oxygenated cofactor may initiate a radical translocation that produces a radical at Tyr27, which then reacts with oxygen directly as opposed to a sMMOH type mechanism where the oxygen inserted is derived from the oxygenated-diiron cofactor (34).

In summary, we have shown that CADD from *C. trachomatis* is a manganese-dependent enzyme that likely employs a dimanganese active site to catalyze two monooxygenase reactions. We provide strong evidence that Tyr27 serves as a sacrificial substrate to serve as a precursor to the aromatic portion of pAB and Lys152 may participate as a sacrificial amino group donor. Future biochemical, spectroscopic, and crystallographic studies will be necessary to reveal the nature of the active cofactor required for the pAB synthase activity of CADD as well as details of the mechanism involved in the self-sacrificing reaction.

## Experimental procedures

Full experimental procedures are provided in the supporting information.

## Acknowledgements

Thank you to Drs. Emily Mevers, Andrew Lowell, and Robert White for insightful discussions and to Eric Truong for help with protein purification. Thank you to the Sobrado Lab (Virginia Tech, Department of Biochemistry), especially Sydney Johnson, for assistance with oxygen consumption and H_2_O_2_ production assays.

## Funding and additional information

This study was supported by start-up funds to KDA (Department of Biochemistry and the College of Agriculture and Life Sciences at Virginia Tech). The Virginia Tech Mass Spectrometry Incubator is funded by the Fralin Life Sciences Institute at Virginia Tech.

## Conflict of Interest

The authors declare that they have no conflicts of interest with the contents of this article.

## Abbreviations

The abbreviations used are:

CADD: chlamydia protein associating with death domains
THF: tetrahydrofolate
pAB: *p*-aminobenzoate
HDO: heme-oxygenase-like diiron oxidase
sMMOH: soluble methane monooxygenase hydroxylase
RNR: ribonucleotide reductase
pHB: *p*-hydroxybenzoate
DTT: dithiothreitol
HQ: hydroquinol.

## Notes

### Competing Interest Statement

The authors have declared no competing interest.

