## Supplemental Text and Figures for "CADD from *Chlamydia trachomatis* is a manganese-dependent oxygenase that employs a self-sacrificing reaction for the synthesis of *p*-aminobenzoate"

### Experimental Procedures

**Materials.** Dithiothreitol (DTT), isopropyl-β-D-thiogalactopyranoside (IPTG), and ampicillin were acquired from GoldBio. NADH, NADPH, EDTA disodium salt, and Miller's Luria Broth were from Research Products International (RPI). All other chemicals and reagents were from MilliporeSigma unless otherwise specified.

**Overexpression and purification of CADD.** The *ct*610 gene cloned into pET19b (CT610_pET19b) with the downstream proteomics predicted start codon was originally obtained from Professor Anthony Maurelli (University of Florida). Site-directed mutagenesis was used to modify one codon to be consistent with the sequence from *C. trachomatis* 434/Bu listed in the NCBI database (Accession: CAP04311) (1).

For overexpression of CADD with an N-terminal 10X-His tag, the CT610_pET19b plasmid was transformed into *E. coli* BL21 and a single colony from a LB/Amp plate was used to inoculate 20 mL of LB broth supplemented with 100 μg/mL ampicillin, which was incubated at 37°C with shaking at 200 rpm overnight. An aliquot (15 mL) of this overnight culture was used to inoculate 1.5 L of LB broth supplemented with 100 μg/mL ampicillin and was incubated at 37°C with shaking at 200 rpm. When the optical density at 600 nm ($OD_{600}$) reached ~0.7, the expression of CADD was induced by the addition of 0.5 mM IPTG. The cells were cultured for 4 hours at 37°C, harvested by centrifugation, and stored at –20°C until the pellet was needed for purification.

A routine purification was performed from 3 L of culture resulting in a cell pellet of ~12 g. The pellet was resuspended in 30 mL 50 mM sodium phosphate, 300 mM NaCl, and 20 mM imidazole (pH 7.4) (buffer A: 20 mM imidazole). The cells were then sonicated on ice and the insoluble cell debris was removed by centrifugation at 15,000 rpm for 45 mins. The cell lysate was loaded into a gravity flow column (1 by 3 cm, ~2 ml of resin) of Ni-nitrilotriacetic acid (Ni-NTA) metal affinity resin (Prometheus), equilibrated with 20 mL buffer A. The column was then washed with 10 mL of buffer A followed by 10 mL of buffer A containing 100 mM imidazole. CADD was then eluted from the column with buffer A containing 500 mM imidazole.

After concentrating the CADD-containing fractions to 2.5 mL using an Amicon centrifuge concentrator (10-kDa cutoff, 15 mL; EMD Millipore), the protein was exchanged into 20 mM Hepes (pH 7.5) using a PD-10 desalting column (Cytiva). The final purified protein in 3.5 mL had an average concentration of ~700 μM monomer (~20 mg/mL). The protein was flash frozen and stored at –80°C until needed for assays. Protein concentrations were determined by the Bradford method with bovine serum albumin as a standard.

***In vitro* enzymatic assays and pAB detection by LC-MS.** Purified CADD described above was thawed on ice. A typical assay was a 500 μL reaction carried out in 20 mM Sodium Hepes (pH 7.5) and contained 154 μM monomer (4.5 mg/mL) protein and 10 mM DTT. Negative control "protein only" reactions contained only 154 μM protein in 20 mM Hepes (pH 7.5). The enzyme reactions were incubated at 37°C for 1 hour before being quenched with 1.5 mL $CH_3CN$. The protein was removed by high-speed centrifugation, the supernatant concentrated under vacuum to 100 μL, and finally analyzed by LC-MS. When performing these assays with anaerobically reconstituted protein (see below), the reconstituted sample was first oxygenated by gently pipetting up and down in an aerobic environment as well as using oxygenated 20 mM Hepes (pH 7.5) to dilute the protein for the reactions. DTT was then added and the reactions proceeded as described above.

For LC-MS analysis, a Waters Acquity TQD mass spectrometer with a Waters Acquity UPLC equipped with an Acquity Premier HSS T3 column (2.1 x 100 mm, 1.8 μm particle size) was used with solvent A as 0.1% formic acid in water and solvent B as 100% methanol. The LC program consisted of 3 min at 95% A followed by a 10-min linear gradient to 50% B at a flow rate of 0.35 ml/min and the injection volume was 10 μl. The MS method was a multiple reaction-monitoring method scanning two pairs: 138.1 and 120.4, 138.1 and 94.4 with a collision energy of 15V and a cone voltage of 30V. The source temperature was 150°C, the desolvation temperature was 500°C, the desolvation gas flow was 800 L/hr, and the cone gas flow was 50 L/hr. MassLynx was used for system operation and data processing.

**Oxygraph detection of CADD oxygenase activity.** Oxygen consumption was measured in a 1 mL reaction using an oxygen electrode system (Hansatech, Amesbury, MA). The protein (200 μL, final concentration of 154 μM or higher) was added to 780 μL of 20 mM Hepes (pH 7.5) and the reaction was initiated by the addition of 20 μL of 0.5 M DTT. The linear portion of the curve was used to determine the oxygen consumption rates for various samples. The oxygen consumption value recorded in the buffer + DTT reaction was subtracted from the oxygen consumption values for reactions containing enzyme to account for the minor background oxygen consumption by DTT alone. The oxygen consumption rate of non-reconstituted CADD was recorded using approximately triple the amount of enzyme as compared to Fe-reconstituted CADD. This was because adding 154 µM of the non-reconstituted protein does not show any measurable oxygenase activity since most of the protein is missing the required diiron cofactor.

**Reconstitution of CADD** A 2.5 mL sample containing no more than 5 mg/mL protein was deoxygenated in a Coy anaerobic chamber (~98% $N_2$/~2% $H_2$) by gently stirring for 2 hours. Then, 2x molar excess ferrous ammonium sulfate was introduced to the sample and gently stirred in the anaerobic chamber for another two hours. Finally, excess Fe was removed using a PD-10 desalting column (Cytiva Life Sciences) eluting the reconstituted protein with deoxygenated 20 mM Hepes (pH 7.5). This sample was capped to keep the sample anaerobic and stored at 4°C overnight or used for *in vitro* enzymatic assays immediately. Experiments showed no significant decrease in activity when the enzyme was left at 4 °C overnight. The Fe-content of purified vs. reconstituted CADD was determined using a colorimetric ferrozine assay (2).

All other metal reconstitutions were performed in the same way as the Fe/Fe reconstitution except for the concentration and type of metal added. For mixed metal reconstitutions (e.g. Mn/Fe), 0.5x molar excess Fe, and 1x molar excess of the other accompanying metal (Mn) was added to the sample and the rest of the reconstitution was carried out described above. For other metal reconstitutions that did not contain Fe (ex: Mn/Mn), 2x molar excess of that metal was added to the sample just as with the Fe/Fe reconstitution. The various metal forms used were: $Cu(II)SO_4$, $Zn(II)SO_4$ and $Mn(II)SO_4$.

**EDTA experiments.** CADD (~5 mg/mL) was gently stirred in the presence of 1 mM EDTA for one hour and then exchanged into 20 mM Hepes (pH 7.5) using a PD-10 desalt column. pAB synthase assays were then performed with the resulting apo-CADD while the remaining portion of the protein was transferred into the anaerobic chamber, deoxygenated with gentle stirring, and reconstituted with Mn, Fe, or Mn/Fe as described above. Finally, these reconstituted protein samples were used to set up final pAB assays.

**$H_2O_2$ production assays.** To analyze $H_2O_2$ production, the Pierce™ Quantitative Peroxide Assay Kit (ThermoFisher Scientific) was used following the manufacturer’s protocol. A 20 µl aliquot from a Fe-reconstituted vs. Mn-reconstituted CADD reaction was removed 30 seconds after initiating the reaction with DTT, mixed with 200 µl of working reagent, incubated at room temperature for 15 minutes followed by measuring the absorbance at 595 nm on a microplate reader. Reactions were performed in triplicate. The amount of $H_2O_2$ present was quantified by comparing to a standard curve.

**$^{18}O_2$ experiments.** Freshly purified CADD was reconstituted with Mn and the sample was used to set up two 1.5 mL pAB synthase assays containing 154 μM CADD. These samples were reduced with 10 mM DTT in the anaerobic chamber, capped and removed from the anaerobic chamber. The $N_2$ and $H_2$ present in the sample vials from the anaerobic chamber were removed under vacuum and one of the vials was flushed with $^{18}O_2$ while the other vial was flushed with $^{16}O_2$ from a syringe filled with air. These reactions were incubated at 37°C in a water bath for 2 hours before the reactions were quenched with 4.5 mL acetonitrile. The protein was removed by high-speed centrifugation and the resulting supernatant concentrated under vacuum to 100 μL for LC-MS analysis on a Waters UPLC connected to a TQD mass spectrometer. The MS method consisted of a MS scan from 50-200 *m/z*, the source temperature was 150°C, the desolvation temperature was 500°C, the desolvation gas flow was 800 L/hr, and the cone gas flow was 50 L/hr.

**LC-MS experiments to identify amino acid modifications.** Mn-reconstituted CADD was used to set up two standard pAB synthase assays in the presence of 10 mM DTT. After incubating these reactions for 1 hour, 25 μg of protein from each sample was diluted to 1 μg/μl in 50 mM triethylammonium bicarbonate, pH 8.5 (TEAB) followed by digestion overnight with 0.5 μg Glu-C endoproteinase MS grade (Thermo Scientific). The digest was diluted in 2: 98 LC-MS grade acetonitrile: water containing 0.1% (v/v) formic acid (solvent A) to 100 ng/μl and 500 ng was analyzed by LC-MS/MS utilizing an Easy-nLC 1200 autosampler/liquid chromatography system, a μPAC compatible nanospray emitter, an Easy Spray nanospray source and a tribid Orbitrap Fusion Lumos mass spectrometer (Thermo Scientific).

The sample was loaded onto an Acclaim PepMap 100 (100 μm x 2 cm, nanoViper C18 5 μm, 100A, Thermo Scientific) serving as a precolumn followed by valve switching to a second Acclaim PepMap 100 (100 μm x 2 cm, nanoViper C18 5 μm, 100A, Thermo Scientific) that served as an analytical column. Flow rate during peptide separation was maintained at 250 nl/min. The LC gradient began at 2: 98 solvent A: solvent B (80: 20 LC-MS grade acetonitrile: water containing 0.1% (v/v) formic acid), increased to 15% B over 2 min then 50% B over 20 min and finally up to 98% B in 1 min with a 2 min hold then back to 2% over 1 min with a 1 min hold at 2% for a total

27 min. Spray voltage was 1800 V, ion transfer temperature was 275˚C, the RF lens was set to 30% and the default charge state was set to 2.

MS data for the m/z range of 400-1500 was collected using the orbitrap at 120000 resolution in positive profile mode with an AGC target of 4.0e5 and a maximum injection time of 50 ms. Peaks were filtered for MS/MS analysis based on having isotopic peak distribution expected of a peptide with an intensity above 2.0e4 and a charge state of 2-5. Peaks were excluded dynamically for 60 seconds after 2 scans with the MS/MS set to be collected at 55% of a chromatographic peak width with an expected peak width (FWHM) of 20 seconds. MS/MS data starting at m/z of 150 was collected using the orbitrap at 15000 resolution in positive centroid mode with an AGC target of 1.0e5 and a maximum injection time of 200 ms. Activation type was HCD stepped from 25% to 35%. For validation of the modification observed at Lys152 in the Glu-C digests, a second run utilized a targeted mass list with the m/z value of 416.23. If a fragment mass of 249.11 was observed in the MS/MS using HCD activation, this triggered a second MS/MS scan utilizing ETD activation with supplemental HCD activation set at 15% using previously calibrated charge-dependent ETD parameters determined using the MRFA 524.3 peak of the Flex Mix tuning mix (Thermo Scientific). Proteome Discoverer v. 2.5 (Thermo Scientific) was used to generate a mascot generic format peak lists which were then searched using Mascot search engine v. 2.7 (Matrix Science) with the following parameters: precursor mass tolerance of ±10ppm, fragment mass tolerance of ±0.1Da, variable modifications of N-terminal Gln to pyro-Glu and oxidation of Met with Glu-C specificity and error-tolerant searching set to true. The databases searched were the common laboratory contaminant protein list included with Proteome Discover and a reference protein for *E. coli* with the recombinant CADD protein sequence with an N-terminal His tag appended.

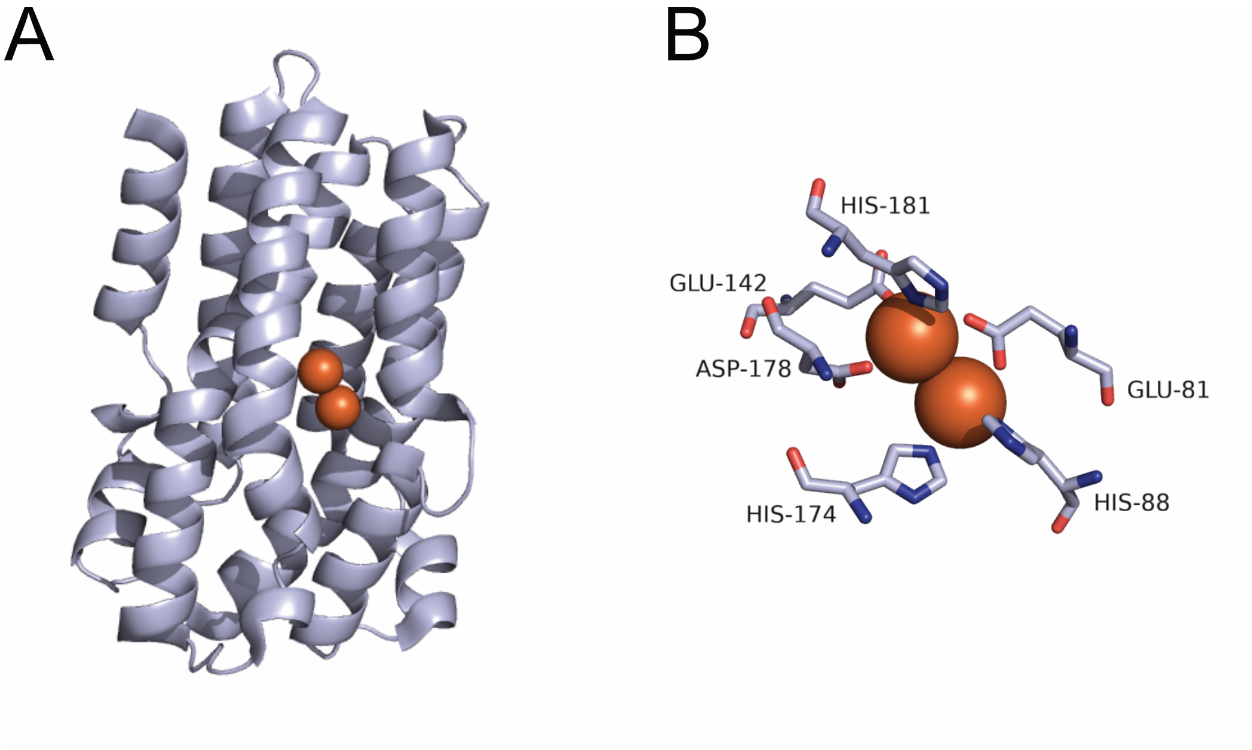


**Figure S1. Crystal structure of CADD.** (A) CADD monomer showing the 7-helix bundle similar to heme oxygenase. (B) Residues coordinating the diiron cofactor. Images were rendered in PyMOL Molecular Graphics System (version 2.5.2) using PDB:1RCW (3).

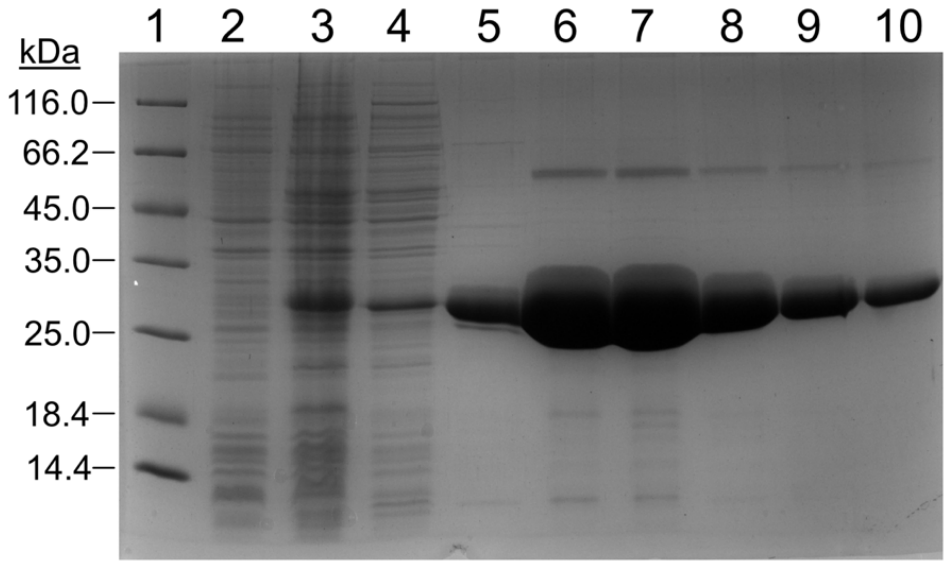


**Figure S2. SDS-PAGE gel of typical his-tagged CADD purification.** Lane 1: molecular weight marker; Lane 2: *E. coli* total cellular proteins before induction with IPTG; Lane 2: *E. coli* total cellular proteins after induction with IPTG; Lane 4: 10% B wash fraction from nickel-affinity purification; Lanes 5-10: 50% and 100% B elution fractions that were pooled and exchanged for subsequent enzyme assays. The monomeric molecular weight of CADD (proteomics predicted start codon) expressed from pET19b with an N-terminal 10x-His tag and enterokinase cleavage site is 29.3 kDa.

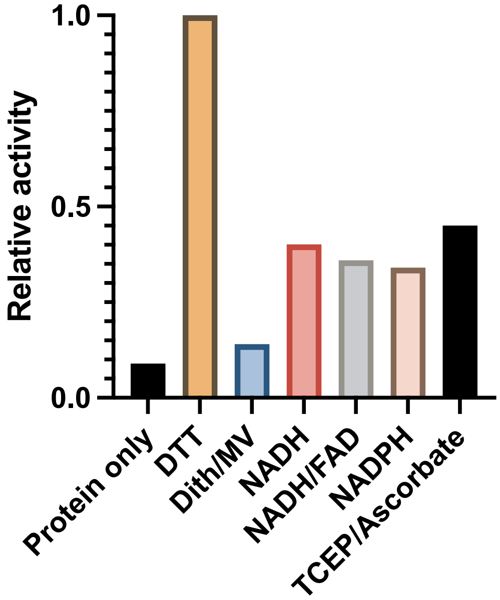


D

**Figure S3. pAB synthase activity of CADD in the presence of various reducing agents.** DTT, dithiothreitol (10 mM); Dith/MV, sodium dithionite (1 mM) + methyl viologen (20 μM); NADH (1 mM); NADH/FAD (both 1 mM); NADPH (1 mM); TCEP, tris(2-carboxyethyl)phosphine (1 mM) + ascorbate (1.  mM).

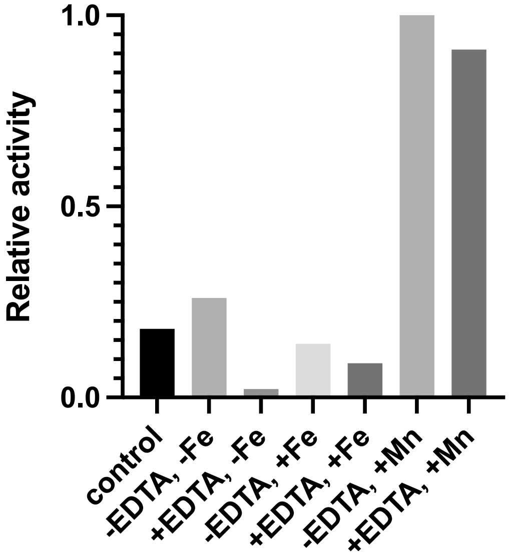


**Figure S4. pAB synthase activity of CADD before and after treatments with EDTA.** All of the samples were from the same pooled protein preps and all of the final reactions contain 154 μM CADD with 10 mM DTT. Control: the typical pAB synthase activity assay in aerobic conditions in the presence of DTT. -EDTA, -Fe: sample was stirred at room temperature in the absence of EDTA, followed by buffer exchange and concentrating before standard pAB synthase assay, this experiment was done to ensure that the stirring and subsequent handling of the protein did not decrease the activity compared to the control. +EDTA, -Fe: stirred in the presence of EDTA followed by buffer exchange and concentrating before standard pAB synthase assay. -EDTA, +Fe: stirred in the absence of EDTA, followed by reconstitution with ferrous ammonium sulfate before pAB synthase activity assay. +EDTA, +Fe: stirred in the presence of EDTA, followed by buffer exchange and reconstitution with ferrous ammonium sulfate before pAB synthase activity assay. -EDTA, +Mn: stirred in the absence of EDTA, followed by reconstitution with manganese sulfate before pAB synthase activity assay. +EDTA, +Mn: stirred in the presence of EDTA, followed by buffer exchange and reconstitution with manganese sulfate before pAB synthase activity assay.

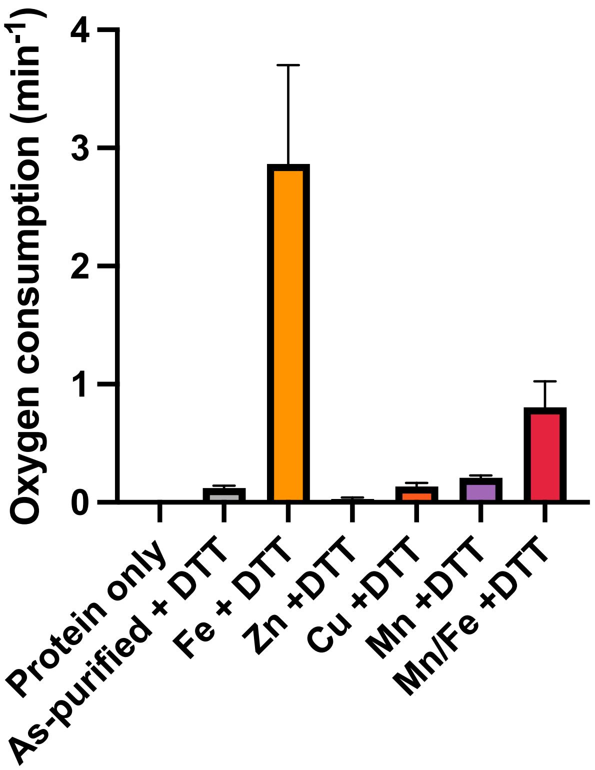


**Figure S5.** Oxygen consumption rate of CADD reconstituted with various metals. Reactions were analyzed using a Hansatech oxygraph with anaerobically reconstituted protein and DTT (10 mM). The minimal non-enzymatic rate of oxygen consumption by DTT in buffer alone was subtracted from the rates of the oxygen consumption observed with protein.

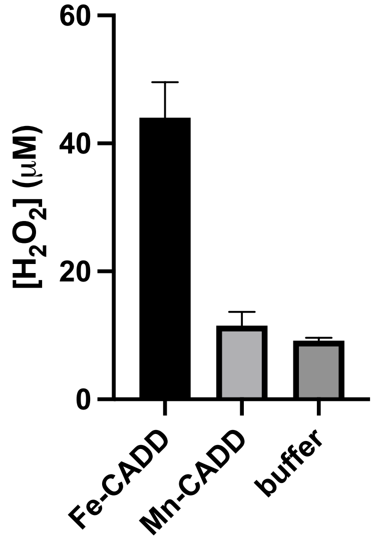


**Figure S6.** Production of $H_2O_2$ by Fe-reconstituted CADD compared to Mn-reconstituted CADD and no protein control. All samples contained 10 mM DTT with 20 mM Hepes (pH = 7.5). Protein samples contained 154 μM protein. $H_2O_2$ production was assayed using a by removing a 20 μl aliquot of the reaction 30 s after initiating the reaction with DTT and employing the Pierce™ Quantitative Peroxide Assay Kit.

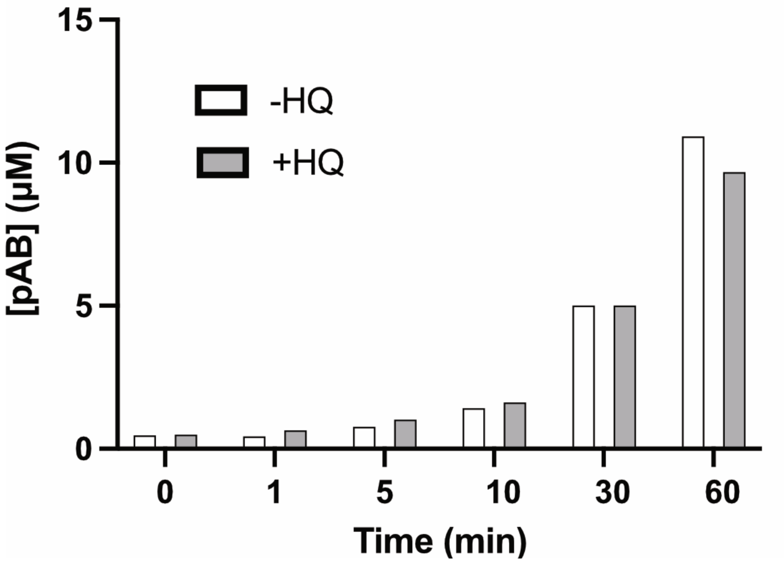


**Figure S7.** Mn-reconstituted CADD pAB synthase activity in the presence vs. absence of hydroquinol (HQ, 0.5 mM).
